# Happiness promotes global processing in haptic perception

**DOI:** 10.1101/2024.07.15.603491

**Authors:** Müge Cavdan, Aycan Kapucu, Katja Doerschner, Knut Drewing

## Abstract

Happy and sad mood promote global and local visual processing, respectively. However, it is unclear whether mood also affects the processing level in haptic perception. Here, we used classical music to induce happy and sad mood in blindfolded participants before they scanned 3D-printed configurations with their fingers. Additionally, we included a neutral group who did not listen to any music. Global shapes were triangles, circles, or squares (33mm) composed of smaller local relief shapes (3 mm): either triangles, circles, or squares. Participants explored a probe stimulus with identical local and global shapes, and two comparison stimuli, matching the probe in local, or global shape. They reported which comparison stimulus appeared more similar to the probe. In the ‘sad’ group, participants chose the locally matching comparison more frequently than in the ‘happy’ and ‘neutral’ groups, revealing that sad mood promotes local processing also in touch. Overall, participants chose the globally matching comparison more often, suggesting that global processing is more prominent in touch than previously assumed.

## Introduction

Feelings modulate how we perceive and interact with the world. Mood modulates cognitive processes and social behavior, e.g., happy feelings promote reliance on accessible general information such as stereotypes (1,2) and scripts (3) while sad feelings have been associated with focusing on details, avoiding stereotypes, which often leads to an improved accuracy of judgements (4). One explanation for this shift of focus is offered by the affect-as-information hypothesis (5,6). According to this theory, mood is a signal that can be used to guide behavior: happy mood signals a low-threat situation that affords global processing whereas sad mood may signal that a situation is potentially threatening and therefor require more detailed processing. An alternative explanation for the association between mood and level of processing suggests that motivation may act as a mediator. According to this idea detailed processing is effortful and may result in a reduction of mood (7). Consequently, when happy, people may be less motivated to perform effortful, detailed processing, in order to maintain their positive mood.

Interestingly, this shift from general, global to detailed, local processing as the mood shifts from happy to sad, has not only been observed in social and cognitive studies but also emerges robustly in visual shape perception (3,8,9). For example, in an experiment by Gasper and Clore (9), ‘happy’ and ‘sad’ participants indicated the similarity of a given shape (triangle or square) that consisted of local elements (triangle or square, also see Kimchi & Palmer (10) to two figures that either matched the global or the local shape of the stimulus. Their results showed that individuals in the sad mood condition more often classified shapes based on local features compared to ‘happy’. In line with this, Curby and colleagues (11) showed that negative mood decreased the holistic processing of faces. The attentional narrowing of attention when mood is low (8,12–16) is interesting, as it appears to ‘override’ the default holistic processing visual shape and scene perception (17,18).

In general, the haptic system applies more weight to local features than global ones when judging shapes (19). Thus, it is not clear how and in what direction changes in mood would modulate the haptic perception of shape. To address these questions, we investigated how positive and negative mood influences the level of haptic shape processing. Upon mood induction, blindfolded participants scanned a haptic shape stimulus with their fingers and indicated which one of the two comparison stimuli was more similar to the probe. Comparisons matched the probe in either global or local shape. A neutral condition with no mood induction (i.e., no music) was included to assess baseline haptic processing. Our working hypothesis was that happy mood would promote global processing while sad mood would promote a focus on local shape features. Our results provide a deeper understanding of to what extent mood-induced changes in attentional focus might be a general perceptual phenomenon.

## Methods Participants

The required sample size was calculated based on a large effect (Cohen’s *d* = 0.8), power of 80%, and alpha 5%. The projected sample size was 42 for one-sided between-subjects *t*-test (G*Power(20)). Accordingly, 42 participants (9 males, *M*_*age*_ = 22.8, age range: 18-30) were randomly assigned to one of the two mood conditions (happy or sad). An additional 21 participants were recruited to the control condition (7 males, *M*_*age*_ = 24.14, age range: 20-34). Eligibility criteria for the study required that participants have no diagnosed mental disorders, as such conditions might influence processing levels (21) and introduce potential confounds in the experiment. They were recruited through the university email system and compensated with 8 €/hour or course credit. None of the participants reported sensory, motor, or cutaneous impairments. The two-point discrimination threshold at the index finger of the dominant hand was 3 mm or better. The study was ethically approved by a local ethics committee LEK FB06 (2017-0034) in accordance with the Declaration of Helsinki(22) excluding the preregistration. Informed consents were obtained prior to the experiment.

### Stimuli and materials

#### Haptic Stimuli

Haptic stimuli were modeled using the OpenSCAD software and 3D printed (Objet30Pro, StratasysLtd., USA) with model photopolymer material (VeroClear) at a resolution of 600 × 600 × 1.600 dpi (x, y, and z axes respectively). As previous research has shown that the haptic system weighs local shape features more than global features(19), we first conducted a pilot experiment where we investigated the detection thresholds of a small triangle, a circle, and a square in order to determine the shape size that would allow us to reduce the natural saliency of the local features relative to the global ones in the main experiment. Stimuli were two-dimensional (2D) triangles, squares, and circles ranging from 1 mm to 10 mm (size: diameter of a circle, one side of a square, and triangle base; shapes embossed with a height of 1 mm on a plane plate). Blindfolded participants touched the shapes and named the shape they perceive. For each shape mean value, the sizes where a person recognized the shapes correctly were calculated. These values were taken as a threshold and then used as the local shape sizes for the later stimuli production in the main experiment.

Nine stimuli consisting of local and global features modelled using OpenSCAD software and 3D printed at a high resolution as in the pilot experiment. The global shapes were printed on a 40 mm (x-axis) × 40 mm (y-axis) long and 5 mm (z-axis) thick base. The stimuli were mounted on a 3D printed plate which has three gaps. This ensured that the stimuli remained fixed on the table during the exploration. Local shape sizes were 3 mm (both x and y-axes) and global shapes size was 33 (both x and y-axes) for square, circle, and triangle with the height of 1 mm (z-axes). The distance between each adjacent local shape was kept constant at 1 mm. The shapes consisted of a circle of circles, a triangle of triangles, and a square of squares as probes while comparison stimuli were a circle of triangles, a square of triangles, a triangle of squares, circle of squares, a square of circles, and a triangle of circles. The stimuli were combined with each other to make triads (six triads in total). Each triad consisted of one probe (i.e., standard) and two comparison stimuli (see Figure 1 for an example). Probes had the same global and local shapes such as square of squares. Comparison stimuli on the other hand, matched either local or global shape of the probe such as square of circles (global match) and circle of squares (local match).

**Figure 1.**
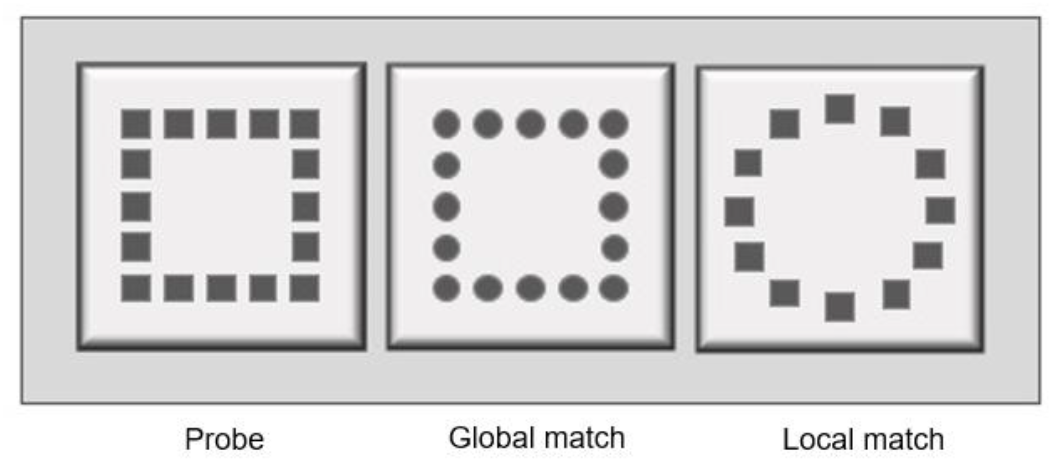
Illustration of an example triad and the plate used in the experiment. Probe: square of squares, global match: circle of squares, local match: squares of circle. The three squares on which the stimuli are placed illustrate the layout of the plate.

#### Mood induction stimuli and manipulation check

Classical music pieces from Ackerley et al. (23) were tested in a pilot experiment in order to affirm their effectiveness in our population^2^. The pieces that evoked strongest happiness and sadness were selected for mood induction in the current study. *Kol Nidrei* by *Max Bruch (1938-1920)* and *Carmen Suite Nr. 1: Les Toréadors* by *Georges Bizet (1838-1875)* were used to induce sad and happy mood, respectively.

In order to measure the efficacy of mood induction, two different scales were used before and after the mood induction, the Self-Assessment Manikin (SAM (24) and the Differential Emotions Scale – DES (25,26)). The SAM measured participants’ valence and arousal levels on a 9-point pictorial depiction (*valence*: 1= very negative to 9= very positive; *arousal*: 1= very calm to 9= very excited). Participants were asked to rate the SAM based on their emotional state. The DES is widely used in psychological research and a reliable method to assess emotions. It consists of several subscales with various adjectives. We selected the sadness and happiness subscales with their corresponding three adjectives per subscale; happiness: *happy, joyful, amused*; sad: *sad, downhearted*, and *blue*. The adjectives were presented with a 9-point scale (0: *not at all*, 8: *extremely*). Participants rated these adjectives based on how much they felt each emotion.

### Procedure

A between-subjects design with two moods – happy and sad – and a neutral condition (no mood manipulation) was used. Participants were randomly assigned to either the happy, the sad mood, and neutral conditions. Upon arrival in the lab, a participant received information about the experiment and signed an informed consent. This was followed by a two-point touch discrimination threshold test to measure any tactile deficiencies (27). Next, participants responded to the mood questions of SAM and DES, presented on a monitor screen, using a numpad. The presentation order of the questionnaires was randomized. After completion of the questionnaires, a classical music piece, either happy or sad - depending on the condition, was played while no music was played in the neutral condition. After this, SAM and DES questionnaires were completed a second time in happy and sad conditions. Participants then performed the Kimchi-Palmer task (10), as described next. In the neutral condition, participants completed the SAM and DES questionnaires a second time after approximately 15-20 minutes had passed.

On each trial, a probe (i.e., same global and local shape) was always presented in the first gap of the mounted plate (Figure 1). The two comparison shapes (one matching the global shape of the probe and the other matching the local shape) were presented in the next two gaps of the mounted plate. The presentation order of the second and third gaps were counterbalanced. Blindfolded participants were asked to explore each stimulus of the triad sequentially (from first gap to third) using lateral movements with the index finger of the dominant hand. Participants were able to re-explore the probe and the comparison stimuli as many times as wanted. Their task was to indicate which of the two comparison stimuli was more similar to the probe.

Each of the six triads was repeated 6 times which resulted in 36 trials in total. The overall procedure lasted 30-45 minutes in happy and sad music conditions while it lasted 20-30 minutes in the neutral condition.

### Data analysis

All analyses were conducted in SPSS 25 software (SPSS inc.) and figures are plotted in MATLAB R2022b (MathWorks inc.).

Cronbach’s alpha was used to calculate reliability per happiness and sadness subscales (DES) for pre- and post-induction in happy and sad conditions and pre- and post-Kimchi-Palmer task in neutral condition. All eight analyses showed good to excellent consistency (Cronbach’s *a* = .70 to .96) allowing for averaging across adjectives. Next, we calculated the average values across the three adjectives per mood (i.e., happiness: happy, joy, and amused; sadness: sad, downhearted, and blue) which were then submitted to an ANOVA to compare happy and sad conditions. Additionally, first mood measurements were submitted to an ANOVA to test whether the baseline mood was comparable across conditions.

For the Kimchi-Palmer task, the frequency of selecting the local comparison shape over the global one was calculated from the raw data of each participant. Selecting the local match indicated that the participant relied on local shape processing. The percentage of local processing across all trials was calculated per participant in order to conduct a one-way analysis of variances (ANOVA) to examine the effect of condition on the level of processing. This was followed by Bonferroni-corrected multiple comparisons to further investigate group differences.

Greenhouse-Geisser correction was applied when the sphericity assumption was violated. Similarly, degrees of freedom were adjusted when Levene’s test indicated unequal variances.

## Results

### Global and local processing

A one-way ANOVA showed a significant main effect of condition: *F*(2, 62) = 7.92, *p* < .001, η^2^ = .21. Specifically, local processing was significantly less frequent after listening to the happy music piece, than after listing to the sad music piece (*M*_*diff*_ = -11.24, *SE* = 4.08, *p* = .023) (Figure 2). Overall, the global match was selected six times more often than the local one (Figure 2). Also, no music condition (*M* = 3.44, *SE* = 1.05) yielded similar local processing frequency as the happy music piece (*M*_*diff*_ = 4.50, *SE* = 4.08, *p* = .82) while it resulted in less frequent than sad music condition (*M*_*diff*_ = -15.74, *SE* = 4.08, *p* < .001).

**Figure 2.**
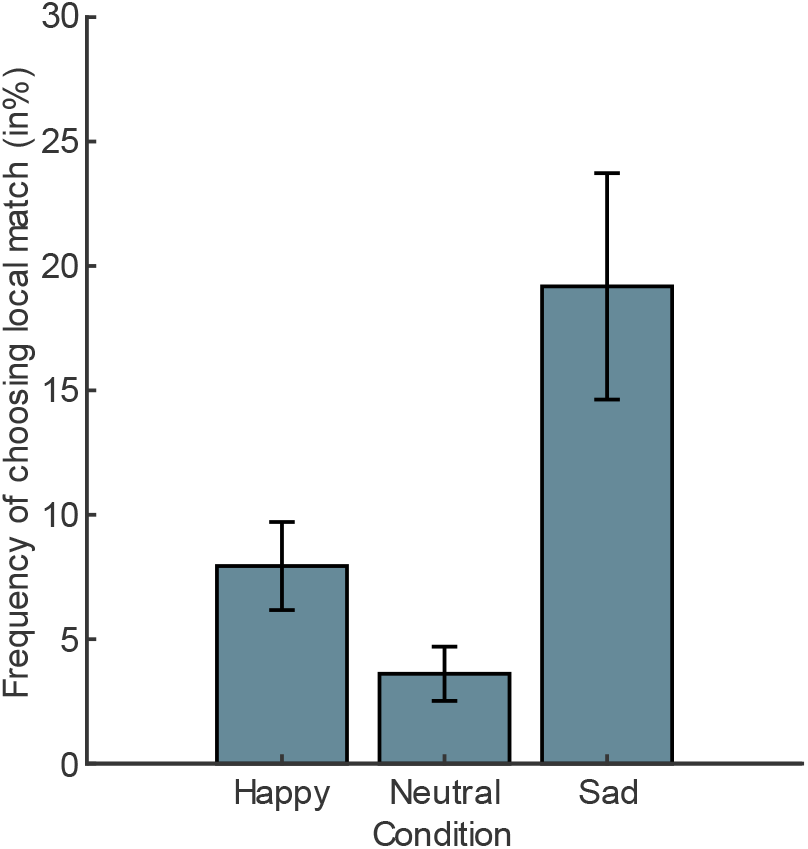
Mean frequency of choosing local math. Error bars correspond to one standard error of the mean.

Next, we tested whether there was an effect of global-local shape combination in level of processing. To this end, we conducted a repeated-measures ANOVA with the variable shape triad (6 levels). The main effect of shape triad was not statistically significant, *F*(4.05, 251.31) = 1.02, *p* = .40, *η*_*p*_^*2*^ = .02, showing that the percentage of local match choice was not dependent on a specific shape-combination. Finally, to test the possible effects of presentation orders on the level of processing, 6 paired-samples t-tests were conducted (i.e., one per shape-combination). None of the t-test were statistically significant showing that there was no effect of presentation order on the percentage of local match (all *p* >.05).

### Mood manipulation check

The efficacy of mood induction was examined using 4 mixed-design ANOVAs to test the effect of music piece (levels: *happy* and *sad*) and time point (levels: *pre-induction* and *post-induction*) separately on mean ratings for Self-assessment Manikin (SAM) valence, SAM arousal, Differential Emotion Scale (DES) happiness, and DES sadness (see Table 1 for the summary of the results).

**Table 1.** SAM Means (with standard deviations) for happiness, sadness, valence, and arousal ratings across happy and sad mood before and after mood induction. Bold numbers in the post test indicate the effect is in the expected direction. * indicates the effect was statistically significant.

| Scales | Happy mood |  | Sad mood |  |
| --- | --- | --- | --- | --- |
|  | pre-test | post-test | pre-test | post-test |
|  | M (SD) | M (SD) | M (SD) | M (SD) |
| Valence (SAM) | 7.00 (1.10) | **7.52** (1.54)* | 7.05 (.80) | **6.48** (1.40)* |
| Arousal(SAM) | 5.52 (2.02) | **6.24** (2.30) | 4.86 (1.71) | **3.67** (1.53)* |
| Happiness(DES) | 5.92 (1.45) | **6.56** (1.14)* | 5.97 (1.03) | **5.51** (1.07)* |
| Sadness(DES) | 2.33 (1.37) | **2.09** (1.22) | 2.13 (1.51) | **2.33** (1.24) |

Additionally, we tested whether the individuals in all conditions had a comparable baseline (i.e., before mood induction) mood with one-way ANOVA on SAM valence, SAM arousal, and DES happiness and sadness subscales. We expected all conditions to have a comparable baseline.

*SAM*. As expected, listening to the happy music piece increased **valence** towards positive, while listening to the sad music piece decreased it: The main effects of mood and time point were not statistically significant for valence, *F*(1,40) = 2.21, *p* = .15, *η*_*p*_^*2*^ = .05, *F*(1,40) = 0.17, *p* = .90, *η*_*p*_^*2*^ = 00, respectively. However, the interaction between time and mood was statistically significant, *F*(1,40) = 8.94, *p* < .01, *η*_*p*_^*2*^ = .18. Follow-up analyses showed that, in the happy mood condition, participants were more positive in the post-test compared to the pre-test, *t*(20) = -2.33, *p* = .015 (one-sided). Also expectedly, after listening to the sad musical piece, participants reported to be in a more negative mood than during the pre-test phase, *t*(20) = 1.98, *p* = .03 (one-sided, Figure 3).

**Figure 3.**
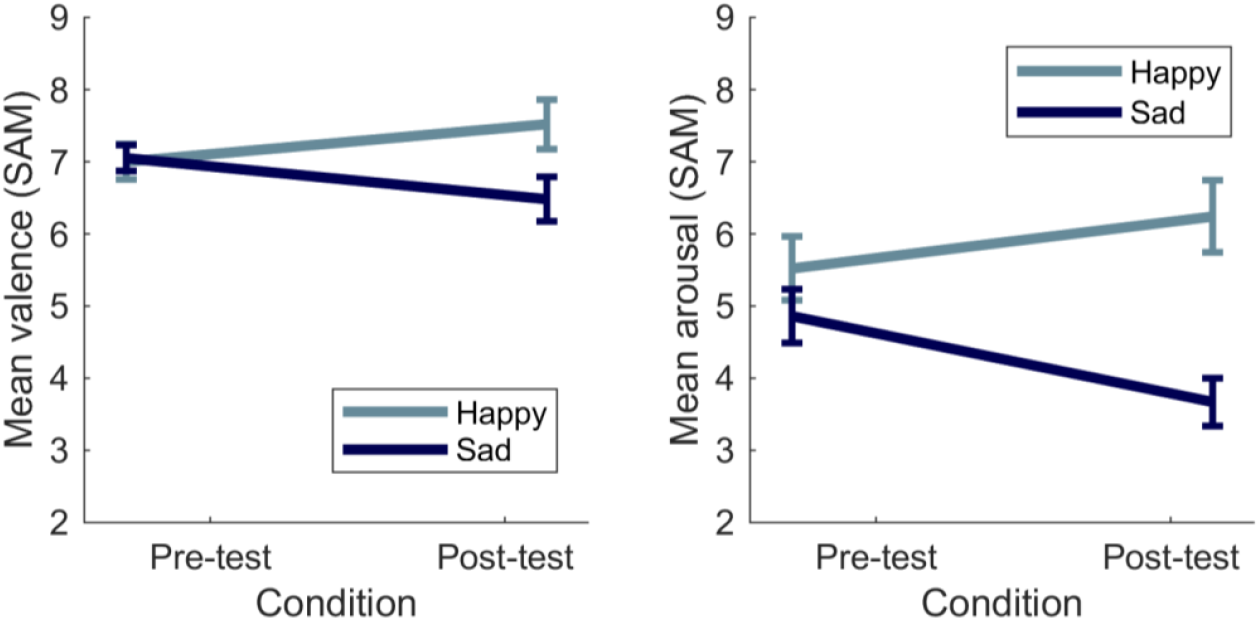
Mean valence (left panel) and arousal (right panel) during pre-test and post-test for happy and sad mood induction conditions. Error bars correspond to one standard error of the mean.

The sad music piece decreased **arousal**, as expected: The main effect of mood, *F*(1,40) = 9.81, *η*_*p*_^*2*^ = .20 and the interaction between time point and mood, *F*(1,40) = 11.24, *η*_*p*_^*2*^ = .22, were statistically significant (both *p* < .01). However, the main effect of time point was not statistically significant for arousal, *F*(1,40) = .70, *p* = .41, *ηp*^*2*^ = .02. Follow-up analyses showed that for happy mood, there was no significant arousal difference between pre-test and post-test, *t*(20) = -1.92, *p* = .07. As expected, for sad mood, arousal decreased on post-test compared to pre-test, *t*(20) = 2.78, *p* < .01 (Figure 3).

Finally and expectedly, prior to listening to the music, there was no difference in valence, *F*(2,60) = .54, *p* = .54, *η*_*p*_^*2*^ = .01 and arousal, *F*(2,60) = 2.31, *p* = .11, *η*_*p*_^*2*^ = .07 across conditions.

*DES*. Participants in the happy mood were happier upon mood induction. However, participants in sad mood did not feel significantly sadder after manipulation which will be discussed further.

For the happiness subscale, neither the main effect of time point, *F*(1, 40) = .28, *η*_*p*_^*2*^ = .01, nor the main effect of mood, *F*(1, 40) = 2.34, *η*_*p*_^*2*^ = .06 were statistically significant. However, the interaction between time point and mood was statistically significant *F*(1, 40) = 11.04, *p* < .01, *η*_*p*_^*2*^ = .06. In the happy mood condition happiness increased between pre- and post-test, whereas in the sad mood condition happiness decreased between pre- and post-test: *t*(20) = -3.11, *p* < .01, *t*(20) = 1.78, *p* < .05 (both one-sided).

For the sadness subscale, although people in the sad mood felt less happy and more sad, this did not reach significance: The main effect of time point: *F*(1, 40) = .01, *η*_*p*_^*2*^ = .00, mood: *F*(1, 40) = .00, *η*_*p*_^*2*^ = .00 and their interaction: *F*(1, 40) = 1.74, *η*_*p*_^*2*^ = .04 (all *p* > 0.05) were not statistically significant.

Additionally, prior to listening to the music pieces, individuals had comparable happiness, *F*(2, 60) = .30, *p* = .75, *η*_*p*_^*2*^ = .01, and sadness, *F*(2, 60) = .15, *p* = .86, *η*_*p*_^*2*^ = .01 levels.

## Discussion

When we look at a scene, we have the ability to focus on a single element or to take in the entire shape formed by these individual elements. Gasper and Clore (9) suggest that whether we perceive the entire shape or the single element depends on our mood. Namely, while happy mood fosters the entire shape view (i.e., global processing), sad mood fosters the single element view (i.e., local processing). Here, we showed that this effect is not limited to vision but a broader mechanism that operates at the perceptual level, and is therefore also present in the haptic modality. Individuals in the happy mood condition matched the probe with the global shape more often than individuals in the sad mood condition. Interestingly, individuals, independent of the mood condition, more often matched the global target to the probe – the global match was selected overall six times more often than the local one. This overall global perception preference is interesting considering that typically people focus on local details when exploring objects with their hands (19,28,29). However, the objects used in these earlier studies were larger than a typical hand, which might have pushed individuals to do sequential exploration, which might have resulted in increased focus on local detail. Thus, the current global precedence finding might be due to the overall size of the global form (30,31), as well as the density (30–32) of the local features (see (33) for a meta-analysis on the environmental factors moderating global precedence. Although local-global focus preferences did not receive the same attention in haptic perception as in vision, a recent haptic study reported a similar global preference as we observed in our experiment: the smaller the local shape the more global processing was observed (31). Therefore, the level of processing seems to critically depend on the number and relative size of the elements (33,34). It would be interesting, in future experiments, to identify the local-global relative size relationship where the perceptual focus switches.

The general observed global preference suggests that the most available and easily accessible information was the global form of our stimuli. The usage of global information, however, was modulated by the mood: Individuals in the happy mood condition selected the global match to the probe more frequently than individuals in the sad mood condition, and this effect was independent of the presentation order or the geometrical features (circle, triangle, square). Our finding is in line with the levels-of-focus hypothesis arguing that individuals in happy mood condition are more likely to perceive the easily accessible knowledge resulting in broader attentional processing (9). In contrast, sad mood would result in increased detailed (local) processing.

Not only mood influences the processing of the stimuli but also using a certain type of processing (global or local) might facilitate a mood (35). This has been also observed for face identification – global processing facilitated happy face identification while local processing facilitated sad face identification (36,37). Future research should investigate the possible bidirectionality between mood and level of processing in haptics.

Our work might have potential clinical relevance. For example, obsessive compulsive individuals tend to focus on details (38,39), a cognitive style, that has been associated with local interference of small details when the task was the identification of global information (40). Future studies should test whether individuals with obsessive compulsive cognitive style use more local processing compared to a control group also in haptic tasks. Insights from such studies potentially yield the development of novel diagnosis tools for this group of patients.

In conclusion, individuals in happy mood are more likely to *feel* the entire shape. Using a stimulus set consisting of local and global features and a haptic shape perception task, we found that happy mood fosters global processing compared to sad mood. To the best of our knowledge, this is the first study that extended the levels-of-processing effect found in vision to the haptic domain. Unlike previous research in haptic perception, we also found global precedence when people perceive different geometrical shapes. Therefore, more research is needed to understand the different components of global and local processing in human haptic perception.

## Author contributions

Conceptualization: M.C., A.K., K.D., K.Dr; Methodology: M.C., K.Dr.; Investigation: M.C.; Data curation: M.C.; Formal analysis: M.C., A.K.; Visualization: M.C., K.D.; Writing— original draft: M.C.; Writing—reviewing and editing: M.C., A.K., K.D., K.Dr; Supervision: M.C., A.K., K.D., K.Dr.; Funding Acquisition: K.D., K.Dr.; Project Administration: M.C., K.Dr.

## Acknowledgement

Authors thank Hatice Dokumaci for the haptic stimuli and data collection, Yeter Nazlican Kara for the data collection, Rochelle Ackerley for sharing the classical music pieces. This research was supported by Deutsche Forschungsgemeinschaft (DFG, German Research Foundation) – project number 222641018 – SFB/TRR 135 (A5 and B8) and EU Marie Curie Initial Training Network “DyVito” (H2020-ITN, Grant Agreement: 765121). K.Dr and M.C. were supported by DFG-project no. 502774891-ORA project “UNTOUCH”, K.Do was supported by Research Cluster “The Adaptive Mind”, funded by the Excellence Program of the Hessian Ministry of Higher Education, Science, Research and the Arts. Artificial intelligence.

## Funding

This research was supported by Deutsche Forschungsgemeinschaft (DFG, German Research Foundation) – project number 222641018 – SFB/TRR 135 (A5 and B8) and EU Marie Curie Initial Training Network “DyVito” (H2020-ITN, Grant Agreement: 765121). K.Dr and M.C. were supported by DFG-project no. 502774891-ORA project “UNTOUCH”, K.Do was supported by Research Cluster “The Adaptive Mind”, funded by the Excellence Program of the Hessian Ministry of Higher Education, Science, Research and the Arts.

## Lead contact

Further information and requests for the resources should be directed to lead contact, Müge Cavdan.

## Data and code availability

Data have been deposited on the Open Science Framework and will be publicly available upon publication. Any additional information required to reanalyze the data reported in this work is available from the lead contact upon request.

## Footnotes

2 Total of 20 participants participated in the between-subjects pilot experiment. Participants were randomly assigned to either happy or sad condition. Participants in the happy condition listened to the happy classical music pieces while participants in the sad condition listened to the sad classical music. Although more than one classical piece was presented in the experiment, we will be focusing on the pieces subject to the current article. Upon listening to either *Carmen Suite Nr*.*1* : *Les Toréadors* or *Kol Nidrei* participants indicated the strongest level of each emotion they experienced while listening to the music piece on a 9-point scale (0: not at all, 8: extremely). The following emotion words were rated in a random order; happiness: *happy, joyful, amused*; sad: *sad, downhearted*, and *blue*. The average scores across happiness and sadness were calculated. Two-independent samples t-test were used to test the efficacy of the classical music pieces. For happiness, the happy music piece yielded a higher rating than the sad piece, *t* (18) = 4.65, *p* < .01, *d* = 2.08. For sadness, the sad music piece yielded a higher rating than the happy piece, *t* (18) = -5.40, *p* < .01, *d* = 2.41. These results clearly show the efficacy of both classical music pieces in inducing the desired target moods.

## Notes

### Competing Interest Statement

The authors have declared no competing interest.

### Summary of Updates

A control condition has been included to the revised form of the manuscript.

